# Gestational exposure to high heat-humidity conditions impairs mouse embryonic development

**DOI:** 10.1101/2024.04.22.590580

**Authors:** Avinchal Manhas, Amritesh Sarkar, Srimonta Gayen

## Abstract

Unprecedented rates of global warming have created an existential challenge for the sustainable survival of species on this planet. Tropical conditions of high ambient temperature and relative humidity are extremely vulnerable to maternal-child health. Emerging epidemiological studies depict that exposure to extremely hot weather conditions during pregnancy leads to an array of adverse pregnancy outcomes, such as low birth weight, stillbirth, preterm delivery, congenital abnormalities and adult-onset disorders. A lack of understanding of the underlying molecular pathophysiology limits us from developing an effective combat strategy in terms of targeted therapeutics to improvise the combined teratogenic effects of high heat-humidity in pregnancy. To address this, it is important to delineate the effect of hot-humid weather conditions on the process of embryogenesis. However, working with human embryos is technically and ethically challenging. In this study, we have established a mouse model of heat-humidity stress during pregnancy, which essentially recapitulates the adverse pregnancy outcomes observed in humans. Importantly, we have profiled the impact of high heat-humidity exposure during gestation at different stages of embryogenesis using this mouse model. Our results indicate that the teratogenicity of heat-humidity stress gets manifested in a cumulative manner, starting from the pre-implantation stage and becomes severe at the mid-gestation, culminating in significantly higher embryonic deaths and malformations at the late gestational stage of mouse embryogenesis. Overall, our study paves the path for exploring the underlying molecular players that get dysregulated under gestational exposure to hot and humid conditions, resulting in severe embryonic defects.

## Introduction

Climate change can be visualized as a long-term shift in global weather patterns, majorly driven by global warming events (*Sixth Assessment Report — IPCC*, n.d.). Currently, global climate change poses a serious threat to the sustainable survival of all species, including humans (Lee & Romero, n.d.; McMichael et al., 2006). The global surface temperature of the Earth has risen by a factor of 1.0-1.5°C since the pre-industrial era, with increased incidence of heatwave events and extremes of humidity in different parts of the globe (Lindsey & Dahman, 2024). This global warming has been reported to have catastrophic impacts on the physiological processes of living organisms (Haines et al., 2014; McCarty et al., 2009). Unlike ectotherms, thermoregulatory adaptive strategies allow endothermic animals to maintain a constant body temperature in a variety of environmental conditions (Bennett & Ruben, 1979; Pörtner, 2004). However, with the current status of global warming, it is challenging even for the endotherms to regulate their internal body temperature at the optimal level. Moreover, high environmental humidity aggravates the perception of heat by compromising essential thermoregulatory functions such as reduced perspiration (Jin et al., 2017). Through the years, mounting evidence indicates a concomitant increase in incidences of cardiovascular illnesses, heatstroke events, mortality, adverse pregnancy outcomes and morbidity (Gasparrini et al., 2015; Kjellstrom et al., 2009; Riley et al., 2018) with rising heat-humidity index. On a global scale, retrospective cohort studies show pregnant women to be the most vulnerable upon exposure to high-temperature conditions, leading to aberrant pregnancy outcomes (Strand et al., 2011).

Prenatal maternal stress is reported to be associated with numerous birth complications and late-onset disorders (Baumgartner and Chrisman, 1987; Cardwell, 2013; Solano et al., 2011; Walsh et al., 2019). In the context of heat as a prenatal stressor, exposure to high temperatures is reported to cause a multitude of pregnancy complications, with both short and long-term implications in post-partum fetal growth in humans (Boni, 2019; Takahashi, 2012). Apart from humans, heat stress during pregnancy has also been reported to cause adverse reproductive outcomes in livestock animals (Bell et al., 1989; Kumar et al., 2021; Skibiel et al., 2018; Thornton et al., 2022; Zhao et al., 2020). Now, this has turned into a critical health emergency for maternal-child health with increased evidence of stillbirths, preterm birth, low fetal birth weight, developmental deformities, congenital defects and even late-onset disorders in infants (Auger et al., 2014, 2017; Chersich et al., 2020; Lajinian et al., 1997; Sun et al., 2019; Wang et al., 2013). This has been a matter of concern for several African and Asian countries, including India, where the socioeconomic conditions demand majority of the workforce to engage in outdoor work (Goyal et al., 2023; Kothawale et al., 2012; Kumar Dash et al., 2022; Nyadanu et al., 2023). As a result, majority of the pregnant women belonging to middle- and low-socioeconomic countries get exposed to high ambient temperature and high relative humidity with less access to medical care during their entirety of gestation (Rekha et al., 2024). However, to date, there is no targeted protection strategy available to combat heat-humidity stress vulnerability during pregnancy (Pillai & Dala, 2023). To address this, we need an extensive investigation into the pathophysiology of heat-humidity exposure during pregnancy, which remains poorly understood. To explore this, it is imperative to understand the impact of high heat-humidity exposure on modulating the trajectory of embryonic development.

However, experimental investigation into the impact of gestational thermal insult on human embryonic development has always been a roadblock due to ethical and technical challenges as well as the non-availability of a surplus number of human embryos. In this context, mice are considered an excellent model system to study human embryogenesis (Morse and Fox, 1981). Therefore, in this study, we have investigated the integrated effect of gestational exposure to elevated temperature associated with high relative humidity on mouse embryonic development. We show that gestational heat-humidity stress (HS) severely impairs embryonic development in mice and thereby hugely impacts gestational outcomes. In fact, we find that our mouse model of gestational HS recapitulates the many aberrant outcomes observed in humans (Basu et al., 2018; McElroy et al., 2022). Moreover, morphometric analysis of developmental defects at different stages of gestation indicated a cumulative nature of the teratogen that initiates at the pre-implantation stages of mouse embryonic development. Together, our HS mouse model paves the path for future investigation into the molecular pathophysiology of heat-humidity stress during mammalian pregnancy. We believe that this model system can be leveraged further to develop effective mitigation strategies for humans in the near future.

## Results

### Heat-humidity stress during pregnancy affects gestational outcomes in mouse

We aimed to explore whether a mouse model of HS during gestation recapitulates the adverse birth outcomes observed in humans. To explore this, we designed our experimental strategy to keep two independent sets of pregnant dams (7-10 weeks old) in two separate conditions, one in control and another under HS. Briefly, one group of pregnant dams were given a 3-hour long exposure (13:00 IST – 16:00 IST) to high temperature and relative humidity (39±1°C, 60±5% RH) daily for the entire period of gestation till gestational day 18.5 (GD18.5) (Fig.1). Simultaneously, we exposed another group of pregnant dams to 25±1°C, 50±1% RH in a similar manner, which served as our experimental control (Fig. 1). All animals belonging to control and experimental groups were kept under similar housing conditions. We compared the gross gestational outcomes between the control vs. HS groups. Interestingly, we observed that exposure to HS during gestation led to a significant reduction in the total number of viable pregnancies. We found that while the live pregnancy rate for the control group was 91% (n=24), it reduced to 48% in the case of the HS group (n=29) (Fig. 2A). Importantly, the average pups delivered per pregnant dam (litter size, LS) was also found to be significantly reduced in the HS group (LS=3.79; n=29) as compared to the control (LS=10.75; n=24); (Fig. 2B). On the other hand, it has been reported that prepartum weight gain in pregnant females positively correlates with litter size, neonate weight at birth and influences various other parameters of pregnancy (Dahake & Shaikh, 2019; Watkins et al., 2008). To investigate this, we comparatively analysed the maternal weight gain status throughout the entire gestational period between the control and HS groups. We found that pregnant females exposed to high heat-humidity conditions were compromised in the expected body weight gain status as compared to the control. Differences in the rate of increase in body mass for control mice vs HS mice peaked off significantly from GD9.5 and increased thereafter, indicative of perturbed pregnancy in the latter case (n=29 for HS, n=24 for control) (Fig. 2C). Interestingly, uteri collected from HS pregnant dams that did not deliver any live pups after GD19.5, revealed either an absence or loss of pregnancy represented by regressed black masses (Figure 2D). Stillbirth events, defined as the delivery of a dead foetus, were also observed in one HS litter on the day of delivery (Fig. 2E). Moreover, high temperature during pregnancy is known to have several negative implications in post-natal health (Edwards et al., 2003; Roos et al., 2021; Walsh et al., 2019). To assess the immediate effect of heat-humidity stress during pregnancy on the F1 generation, we took a comparative approach to investigate the neonate weight at birth between the control and HS groups. We observed that the HS cohort gave birth to pups with significantly lower mean birth weight (1.28; n=6) as compared to the control F1 pups (1.44; n=11); (Fig. 2F). Altogether, we conclude that heat-humidity stress during mouse gestation affects the gross gestational outcomes characterized by lower rates of viable pregnancy, reduced litter size, low birth weight of neonates, and stillbirth events.

**Figure 1:**
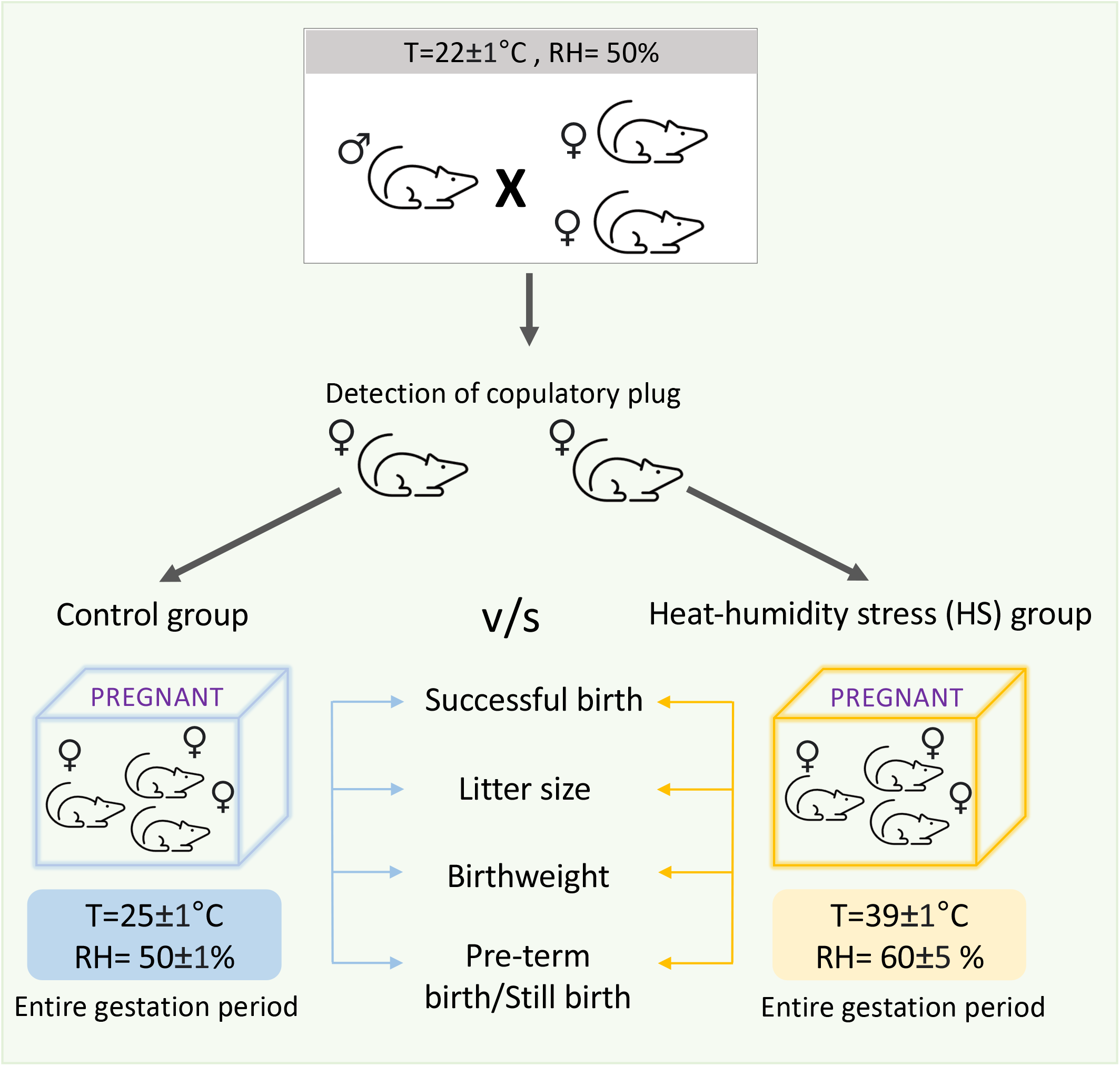
Experimental strategy for profiling gestational outcome upon high heat humidity exposure. Schematic showing the outline of the experimental strategy for gestational exposure to high heat-humidity stress using mouse model. Pregnant female mice were exposed 3hr/day to high temperature (39±1°C) and high relative humidity conditions (60 ± 5%). For the control group, pregnant female mice were exposed to standard temperature 25±1°C and relative humidity conditions of 50±1%. Following their gestational period, gestational outcomes were assessed as shown.

**Figure 2:**
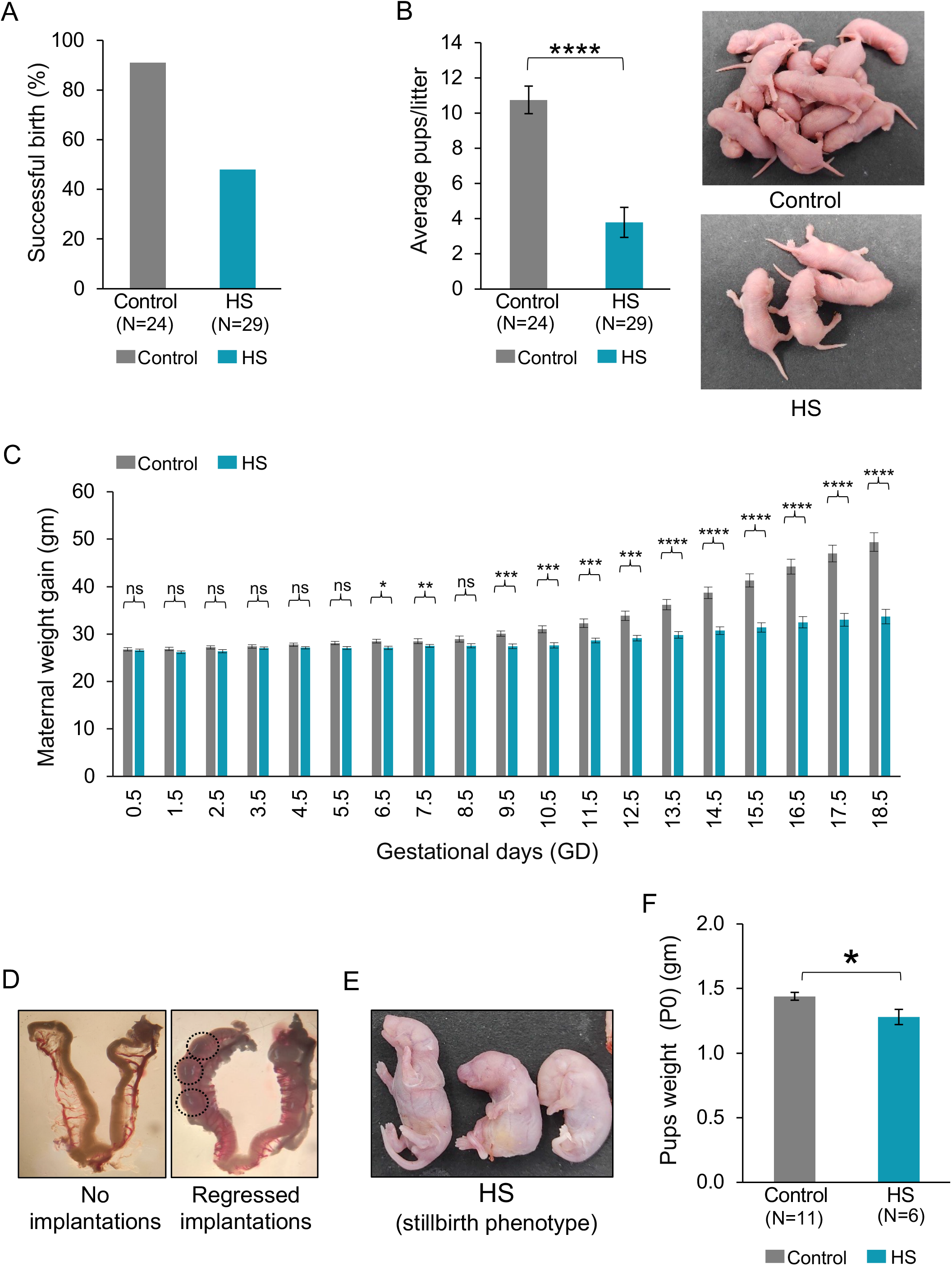
Gestational outcomes upon high heat-humidity stress during mouse pregnancy. (A) Bar plot depicting percentage (%) of pregnant dams giving successful birth after 20 days of gestation (n=24 for control, n=29 for HS). The event of successful birth was attributed to dams that gave birth to live or dead fetuses, irrespectively. (B) Left: Bar plot showing the mean number of pups/dam (litter size) in control (n=24) vs HS group (n=29). (Mann-Whitney U test). Right: Representative images depicting post-partum litter size obtained from the control (up) and HS group (down). (C) Plot showing comparative maternal body weight gain of control (n=24) vs. HS (n=29) group measured between GD0.5 to GD18.5 (GD: Gestational day; Student’s t-test). (D) Representative image showing gross morphology of primigravid uteri at GD19.5 from HS pregnant dams, which did not give birth. Left: No signs of implantations observed. Right: Black-red mass observed, indicative of early post-implantation regression of embryo, indicated in black-dotted circle. (E) Representative images showing still-birth phenotype in HS group. (F) Bar plot showing mean birth weight of p0 pups in control vs HS group (n=11 for control, n=6 for HS) (Student’s t-test). Data are presented as mean n±SEM. *\*p<0*.*05; **p<0*.*01; ***p<0*.*001; ****p<0*.*0001; ns: non-significant*.

### Gestational heat-humidity stress leads to severe defects in embryonic development from early-midgestation and onwards

Next, we aimed to delineate the impact of high heat-humidity exposure on fetal development at different stages of embryogenesis. To explore this, we analysed embryonic phenotypes across five gestational stages – pre-implantation E3.5, post-implantation E8.5, early-midgestation E10.5, mid-gestation E13.5, and at late-gestation E18.5 in control vs. HS group (Fig. 3A). To begin with, we did a comparative analysis of E3.5 blastocysts (pre-implantation) isolated from both control and HS female dams (n=5). Interestingly, we observed a slightly delayed development of the HS blastocysts with signs of improper cavitation (Fig. 3B). However, there were no significant differences in the average number of blastocysts between HS (12.6) and control (13.2) (Fig. 3B). Next, we explored the status of post-implantation embryos at E8.5 stage. We did not find much difference both in morphology and in the mean number of E8.5 embryos between the control (12.33) vs. HS group (7) (n=9) (Fig. 3C). Notably, we observed three HS dams with no uterine implantations upon dissecting at E8.5, compared to one in control.

**Figure 3:**
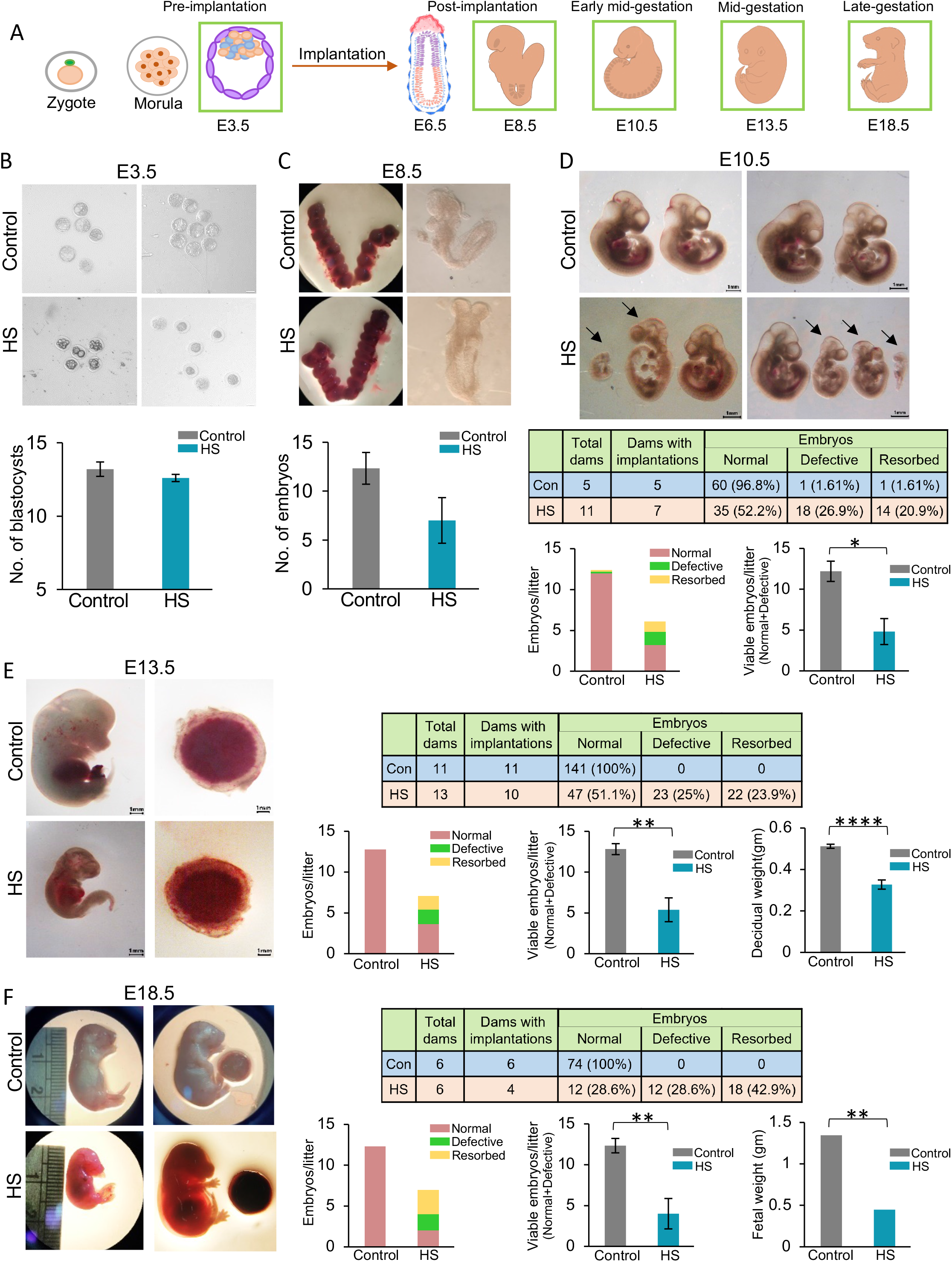
Profiling the impact of gestational exposure to high heat-humidity on mouse embryogenesis. (A) Schematic showing different stages of mouse embryogenesis. Embryonic stages highlighted in green were processed for our analysis. (B) Top: Representative images of E3.5 embryos of HS and control group, scale bar: 50µm. Bottom: Plots showing the quantification of number of E3.5 blastocysts in control vs HS (n=5) group dams. (ns, Welch’s t-test). (C) Representative images and quantification of number of E8.5 embryos obtained from the control vs HS group. (n=9) (ns, Mann-Whitney U test). (D) Top: Representative images of E10.5 embryos. Defective embryos are marked using black arrows; scale bar: 1mm. Bottom: Table showing the quantification of the different categories of embryos (normal, defective, resorbed) in control (n=5) and HS group (n=11). Bar plots depicting the (left) number of different categories of embryos (normal, defective, resorbed) per dam and (right) average number of viable embryos (normal+defective) in control (n=5) and HS group (n=11) (Mann-Whitney U test). (E) Representative images showing control vs HS embryos and placenta of E13.5, scale bar: 1mm. Table showing the quantification of different categories of embryos (normal, defective, resorbed) in control (n=11) and HS group (n=13). Bar plots depicting the number of different categories of embryos (normal, defective, resorbed) per dam (left) and the mean number of viable embryos (normal+defective) (middle) (Mann-Whitney U test) per dam in control (n=11) and HS group (n=13). (Right) Quantification of decidual weight in the control (n=11) and HS group (n=10) (Welch’s t-test). (F) Representative images (E18..5) of embryos and placenta. Table showing the quantification of different categories of embryos (normal, defective, resorbed) in control and HS group. Bar plots depicting the number of different categories of embryos (normal, defective, resorbed) per dam (left) and the mean nuber of viable embryos (normal+defective) (middle) (Student’s t-test) in the control and HS group (n=6). (Right) Quantification of fetal weight in the control (n=6) and HS group (n=4) (Welch’s t-test). Data are presented as mean±SEM. *\*p<0*.*05; **p<0*.*01; ***p<0*.*001; ****p<0*.*0001; ns: non-significant*.

**Figure 4:**
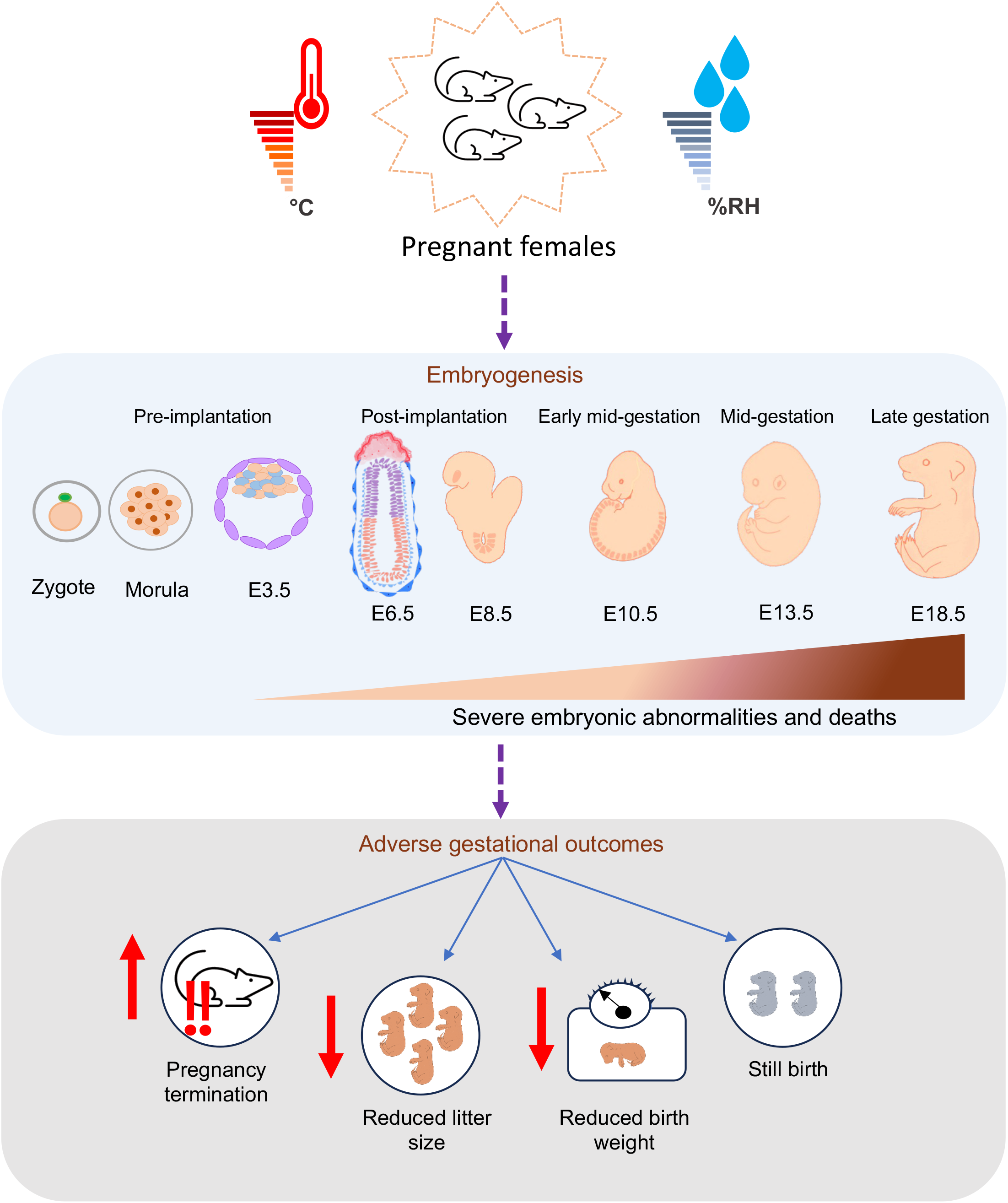
Schematic depicting the effect of heat-humidity stress exposure during pregnancy. Model diagram illustrating the teratogenic effects of exposure to high temperature and high relative humidity during mouse pregnancy. In brief, gestational heat-humidity stress severely perturbs mouse embryonic development in a cumulative manner, with prominent morphological aberration and mortality initiating at the early mid-gestating E10.5 stage. Adverse embryonic phenotypes subsequently get translated into aberrant birth outcomes in the from of loss of pregnancy, reduced neonatal birth weight, stillbirth events and reduced litter size.

Subsequently, we profiled the comparative embryonic phenotype at the early-midgestation E10.5 stage. Notably, while all 5 dams in the control group had visible implantations, only 7 out of 11 dams in the HS cohort had visible implantations, indicating probable loss of pregnancy due to heat-humidity stress. In addition to the loss of pregnancy, we observed severe embryonic defects and mortality in the HS group. We observed many deciduas with completely black and resorbed masses in the uteri of HS dams, indicating “resorbed/dead” embryonic structures (% resorption = 20.9 in HS (n=11) vs. 1.61 in control (n=5)) (Fig. 3D). Moreover, many embryos of the HS group were found to be significantly smaller in size with morphological abnormalities, which we categorised as “defective” (% defective embryos = 26.9 in HS vs. 1.61 in control) (Fig. 3D). Indeed, the overall mean viable embryos (comprising the normal and defective embryos) per dam in the HS group (4.82) was significantly reduced compared to the control group (12.2) (Fig. 3D). Therefore, this finding suggests HS to cause embryonic lethality and malformations observed at the early-midgestation stage of mouse embryogenesis.

Next, we assessed the impact of gestational HS on fetal development in midgestating E13.5 embryos. Here, all 11 control dams had successful implantations in their uteri. On the other hand, only 10 out of 13 dams in the HS cohort had visible implantations at E13.5, which suggested probable loss of pregnancy. As expected, we observed higher resorption frequencies in the HS group (23.9%) compared to none in control (Fig. 3E). Also, in the pregnant dams, the mean decidual weight was slightly reduced in the HS group (0.33g) as compared to control (0.51g) (Fig. 3E). Besides, about 25% of the HS embryos were morphologically quite underdeveloped and retarded in contrast to none in control dams. As a result, the mean viable embryos/dam in the HS group was significantly reduced (5.38) compared to the control (12.82), indicating the death of many embryos by the E13.5 stage (Fig. 3E). Therefore, HS caused severe embryonic defects and deaths observed at the midgestational stage (E13.5) of mouse embryogenesis.

Finally, we assessed the status of embryonic defects upon gestational exposure to HS at the late-gestating E18.5 stage. We observed that embryos from the HS dams were severely retarded as compared to the control. Notably, only 4 out of 6 dams in our HS cohort had visible implantations at E18.5, while all 6 control dams had successful implantations. Unlike control (none), we observed nearly 42.9% of total HS E18.5 embryos to be completely resorbed and appear as dark black masses in the uteri (n=6). Strikingly, 28.6% of the total HS embryos had severe growth defects. On the other hand, all of the control embryos were morphologically normal. Though, we did find another 28.6% of the HS embryos (1.6cm) to be morphologically normal looking, they were also quite smaller in size compared to control (2.5cm) and had improper skin development (Fig. 3F). This data correlates with the significantly reduced average fetal (E18.5) weight in HS group (0.44g) compared to control (1.34g) (Fig. 3F). Taken together, our morphometric analysis suggests that exposure to high heat-humidity during pregnancy hampers murine embryonic development dramatically. Cumulatively, we observed high frequency of pregnancy loss events post-implantation (33.3% in E8.5; 36.4% in E10.5; 23.1% in E13.5; 33.3% in E18.5), which strongly suggests the event of implantation to be extremely vulnerable to high temperature and humidity. Moreover, we find that visible phenotypic aberrations get aggravated in a cumulative manner, starting from early-midgestation at the E10.5 stage, and the defects culminate till parturition.

## Discussion

Though climate change has gained recent attention in the socio-political stage, any attempt to develop sustainable strategies against the catastrophic effects of global warming on the reproductive outcome of mammals have been neglected. Emerging reports suggest deterioration of maternal-child health in recent years, attributable majorly to worsening climate change related to global warming conditions (Zhang et al., 2017).

In this study, we have developed a mouse model of heat-humidity stress during gestation, which recapitulates adverse pregnancy outcomes as reported in humans. As our experimental strategy, ee subjected pregnant female mice to a daily exposure (3 hours/day) of high temperature and humidity (39±1°C, 60±5% RH) for the entire gestational period to mimic conditions as often experienced in the external environment, especially in the tropics. In this mouse model, we find aberrant pregnancy outcomes upon heat-humidity exposure, often characterized by post-implantation loss of pregnancy, fetal growth restrictions, compromised maternal weight gain, low litter size and few stillbirth events. We conclude that our mouse model of gestational heat humidity stress can be leveraged to explore the pathophysiology of heat-humidity vulnerability during pregnancy.

On the other hand, to date, there has been a complete lack of in-depth investigation into the impact of gestational heat-humidity exposure at different stages of embryogenesis. In this study, we leveraged our mouse model of gestational heat-humidity stress to explore the combined effect of high heat-humidity on fetal development, profiled across different developmental stages. Interestingly, we found severe developmental abnormalities and embryonic lethality upon gestational heat-humidity exposure, mostly starting from the early-midgestation stage (E10.5) of mouse embryonic development (Fig 3). Importantly, we observed that the aberrant effect on embryonic development worsened as it moved forward in the developmental time point, indicating high heat and relative humidity to have a teratogenic effect. In contrary to a continual model system of heat stress, a recent study compared the effect of the stage-specific heat exposure and indeed found that it affects mostly at the late gestational phase in mouse (Mayvaneh et al., 2020). However, one recent study did not observe much effect of continual gestational heat stress on mouse pregnancy (Du et al., 2023). Separately, we also found a significant proportion of dams in the HS cohort to have no sign of implantations in their uteri at post-implantation stages, while there was not much differential representation at the pre-implantation stage between HS and control group (Fig 3). We speculate that heat-humidity stress impacts the implantation process and thereby leads to loss of pregnancy in some cases. However, more in-depth investigations are required to disentangle the implantation failures vs. false plug events. On the other hand, we have also observed slight delayed development in pre-implantation blastocyst (Fig 3B). Indeed, few previous reports implicated that transient heat exposure during the pre-implantation development in mice impacts blastocyst development and implantation defects (Bellve, 1972; Edwards et al., 2003; Tian et al., 2013). Altogether, we believe that our study was able to capture the human phenotype strongly due to stringency in experimental designing, simulating the exact environmental parameters found in middle or low-income countries from the tropics.

Overall, our study demonstrates the detrimental impact of high ambient temperature associated with high relative humidity on mammalian pregnancy through mouse model system. At a broader scale, we conclude that global warming associated with elevated heat-humidity index impairs proper embryonic development. Though this study essentially showed the catastrophic effect of HS at different stages of mouse embryogenesis, the underlying molecular pathways that get haywire during the process still remain elusive. Into this, environmental modulations are known to alter the epigenome in a variety of ways. We hypothesize that gestational heat-humidity stress might impact the crucial epigenetic and transcriptional landscape essential during development. Thereby, we need more in-depth investigations in the future at a transcriptomic and epigenomic scale to get a detailed molecular roadmap that gets dysregulated during the process. This will be absolutely crucial to develop targeted therapeutics to be implicated at the human level as an efficient mitigation strategy in the near future.

## Materials and Methods

### Experimental Animals

CD1 mice strain used in this study were obtained from the Central Animal Facility (Indian Institute of Science, Bangalore, India). The entire experiment involving live animals was approved by the Institutional Animal Ethics Committee (approval ID - CAF/Ethics/948/2023). Mice were maintained in standard housing conditions of 22±1°C and 50% relative humidity followed by 12-hour dark/light cycle. All animals were supplied with food and water *ad libitum*. For our experiment, 7-10 weeks old virgin CD1 females were mated with 7-12 weeks old CD1 proven males. Mating was allowed for a maximum period of 5 days. Each morning, female dams were examined for the presence of vaginal copulatory plug, indicating successful mating. The day of identification of a copulatory plug was marked as embryonic day 0.5 (E0.5). After detection of a successful plug, females were randomly assigned to either the experimental group or the control group as described below. Animals were anaesthetized prior to sacrifice followed by cervical dislocation. All efforts were made to minimize suffering.

### Heat-humidity exposure

We have used two separate incubators for our study. The control incubator was set to maintain a temperature of 25±1°C and a relative humidity of 50±1%. Whereas the experimental (HS) incubator was set to maintain a temperature of 39±1°C and a relative humidity of 60±5%. Temperature and humidity were measured using digital thermo-hygrometers.

Animals from both groups (control and HS) were placed inside respective incubators for 3 hours (13:00 IST-16:00 IST) every day and maintained in standard housing conditions. The process was repeated for the entire gestational period or till intermediate gestational stages according to the experimental design. During the entire experiment, body weight gain and food consumption estimates for the pregnant F0 dams were measured starting from Gestational Day – 0.5 (GD0.5). For the F1 generation, the litter size and weight at birth were recorded.

### Pre-implantation (E3.5) embryo isolation

For isolating E3.5 pre-implantation embryos, pregnant dams from each group (control and heat stress) were anesthetized and sacrificed by cervical dislocation. In brief, the uterus was removed and dissected in two halves along the uterine junction. The excess fat was trimmed. The blastocysts were flushed onto a 60mm dish by injecting 2ml of ice-cold DMEM (HiMedia Laboratories, Catalogue #AL007A) using a 27-gauge syringe into the uterine horns under a stereo microscope. Recovered blastocysts were transferred in ice-cold in DBPS with 10% BSA.

### E8.5, E10.5, E13.5, E18.5 embryo isolation

For all post-implantation (E8.5, E10.5, E13.5, E18.5) embryo isolation, the uterus was dissected out and transferred to a petri-plate; washed with ice-cold DPBS twice. The number of mouse embryo implantations was noted and images captured. Decidua were opened out using fine forceps and kept in a 60mm dish containing ice-cold DPBS. For E13.5 and E18.5 stage, individual decidua was weighed individually, and data was recorded. The embryo was gently isolated and separated from the extraembryonic structures. It was picked up and transferred to a new petri dish containing ice-cold 10% BSA in DBPS. For E18.5 stage, both embryo and placenta were weighed separately, and data recorded. Embryos of all the stages were imaged. Fetal resorptions or developmental defects were noted, if any. Linear size was measured for some representative E18.5 embryos for both control and HS group.

### Image acquisition and processing

All E3.5 blastocyst images were captured using Olympus IX73 Research Inverted microscope and cellSens [Ver.2.1] Life Science imaging software with a 10X objective. All other microscopic images were taken under stereo microscope (Olympus SZH-ILLK) using the COSLAB CosLabView software, version: x64, 4.12.24390.20240108.

### Statistical analysis

All data was examined for normality and homogeneity of variance. Normality was tested using the Shapiro-Wilk test and homogeneity of variance was tested using Levene’s test. Student’s t-test was performed for data showing normal distribution and homogeneity of variance. Welch’s t-test was conducted when data followed a normal distribution but did not obey the homogeneity of variance, Mann-Whitney U test was performed when data was not normally distributed.

## Author’s Contribution

SG conceptualised, supervised and acquired the funding for the study. AM and AS performed all the experiments and contributed to data analysis, conceptualisation and writing of the manuscript. SG, AM and AS edited and proofread the manuscript.

## Acknowledgements

This study is supported by Department of Biotechnology (DBT), India (BT/PR30399/BRB/10/1746/2018), Department of Science and Technology (DST-SERB) (CRG/2019/003067), DBT-Ramalingaswamy fellowship (BT/RLF/Re-entry/05/2016) and Infosys Young Investigator award to SG. We would like to acknowledge the Central Animal Facility (CAF), Indian Institute of Science, Bangalore. A.M. acknowledges the financial support received from University Grants Commission (UGC), Ministry of Education, Government of India. A.S. acknowledges the financial support received from Indian Institute of Science, Bangalore (IISc), India.

## References

Auger, N., Fraser, W. D., Sauve, R., Bilodeau-Bertrand, M., & Kosatsky, T. (2017). Risk of Congenital Heart Defects after Ambient Heat Exposure Early in Pregnancy. Environmental Health Perspectives, 125(1), 8–14. 10.1289/EHP171

Auger, N., Naimi, A. I., Smargiassi, A., Lo, E., & Kosatsky, T. (2014). Extreme heat and risk of early delivery among preterm and term pregnancies. Epidemiology (Cambridge, Mass.), 25(3), 344–350. 10.1097/EDE.0000000000000074

Basu, R., Rau, R., Pearson, D., & Malig, B. (2018). Temperature and Term Low Birth Weight in California. American Journal of Epidemiology, 187(11), 2306–2314. 10.1093/AJE/KWY116

Bell, A. W., McBride, B. W., Slepetis, R., Early, R. J., & Currie, W. B. (1989). Chronic heat stress and prenatal development in sheep: I. Conceptus growth and maternal plasma hormones and metabolites. Journal of Animal Science, 67(12), 3289–3299. 10.2527/JAS1989.67123289X

Bennett, A. F., & Ruben, J. A. (1979). Endothermy and Activity in Vertebrates. Science, 206(4419), 649–654. 10.1126/SCIENCE.493968

Boni, R. (2019). Heat stress, a serious threat to reproductive function in animals and humans. Molecular Reproduction and Development, 86(10), 1307–1323. 10.1002/mrd.23123

Cardwell, M. S. (2013). Stress: pregnancy considerations. Obstetrical & Gynecological Survey, 68(2), 119–129. 10.1097/OGX.0B013E31827F2481

Chersich, M. F., Pham, M. D., Areal, A., Haghighi, M. M., Manyuchi, A., Swift, C. P., Wernecke, B., Robinson, M., Hetem, R., Boeckmann, M., & Hajat, S. (2020). Associations between high temperatures in pregnancy and risk of preterm birth, low birth weight, and stillbirths: systematic review and meta-analysis. BMJ (Clinical Research Ed.), 371. 10.1136/BMJ.M3811

Dahake, S. T., & Shaikh, U. A. (2019). A study to assess correlation between maternal weight gain and fetal outcome among primigravidae registered in antenatal clinics. Journal of Family Medicine and Primary Care, 8(11), 3554. 10.4103/JFMPC.JFMPC_756_19

Du, S., Tian, X., Shang, G., Liu, Z., Niu, H., & Ma, J. (2023). Effect of maternal heat stress during pregnancy on reproductive functions of offspring in Balb/c mice. 10.21203/RS.3.RS-2729996/V1

Edwards, M. J., Saunders, R. D., & Shiota, K. (2003). Effects of heat on embryos and foetuses. International Journal of Hyperthermia : The Official Journal of European Society for Hyperthermic Oncology, North American Hyperthermia Group, 19(3), 295–324. 10.1080/0265673021000039628

Gasparrini, A., Guo, Y., & Hashizume, M. (2015). Mortality risk attributable to high and low ambient temperature: a multicountry observational study. The Lancet, 386(9991), 369–375. 10.1016/S0140-6736(14)62114-0

Goyal, M. K., Singh, S., & Jain, V. (2023). Heat waves characteristics intensification across Indian smart cities. Scientific Reports 2023 13:1, 13(1), 1–16. 10.1038/s41598-023-41968-8

Haines, A., Ebi, K. L., Smith, K. R., & Woodward, A. (2014). Health risks of climate change: act now or pay later. The Lancet, 384(9948), 1073–1075. 10.1016/S0140-6736(14)61659-7

Jin, L., Zhang, Y., & Zhang, Z. (2017). Human responses to high humidity in elevated temperatures for people in hot-humid climates. Building and Environment, 114, 257–266. 10.1016/J.BUILDENV.2016.12.028

Kjellstrom, T., Holmer, I., & Lemke, B. (2009). Workplace heat stress, health and productivity – an increasing challenge for low and middle-income countries during climate change. Global Health Action, 2(1). 10.3402/GHA.V2I0.2047

Kothawale, D. R., Kumar, K. K., & Srinivasan, G. (2012). Spatial asymmetry of temperature trends over India and possible role of aerosols. Theoretical and Applied Climatology, 110(1–2), 263–280. 10.1007/s00704-012-0628-8

Kumar Dash, S., Jenamani, R. K., & Mohapatra, M. (2022). India’s prolonged heatwave linked to record poor summer rains. Nature India 2022. 10.1038/d44151-022-00054-0

Kumar, M., Ratwan, P., Dahiya, S. P., & Nehra, A. K. (2021). Climate change and heat stress: Impact on production, reproduction and growth performance of poultry and its mitigation using genetic strategies. Journal of Thermal Biology, 97. 10.1016/J.JTHERBIO.2021.102867

Lajinian, S., Hudson, S., Applewhite, L., Feldman, J., & Minkoff, H. L. (1997). An association between the heat-humidity index and preterm labor and delivery: a preliminary analysis. American Journal of Public Health, 87(7), 1205–1207. 10.2105/AJPH.87.7.1205

Lee, H., & Romero, J. (n.d.). Climate Change 2023 Synthesis Report IPCC, 2023: Sections. In: Climate Change 2023: Synthesis Report. Contribution of Working Groups I, II and III to the Sixth Assessment Report of the Intergovernmental Panel on Climate Change [Core Writing Team. 35–115. 10.59327/IPCC/AR6-9789291691647

Lindsey, R., & Dahman, L. (2024, January 18). Climate Change: Global Temperature. Climate.Gov.

McCarty, J. P., Wolfenbarger, L. L., & Wilson, J. A. (2009). Biological Impacts of Climate Change. In eLS. Wiley. 10.1002/9780470015902.a0020480

McElroy, S., Ilango, S., Dimitrova, A., Gershunov, A., & Benmarhnia, T. (2022). Extreme heat, preterm birth, and stillbirth: A global analysis across 14 lower-middle income countries. Environment International, 158, 106902. 10.1016/j.envint.2021.106902

McMichael, A. J., Woodruff, R. E., & Hales, S. (2006). Climate change and human health: present and future risks. The Lancet, 367(9513), 859–869. 10.1016/S0140-6736(06)68079-3

Nyadanu, S. D., Tessema, G. A., Mullins, B., Kumi-Boateng, B., Ofosu, A. A., & Pereira, G. (2023). Prenatal exposure to long-term heat stress and stillbirth in Ghana: A within-space time-series analysis. Environmental Research, 222. 10.1016/J.ENVRES.2023.115385

Pillai, A. V., & Dala, T. (2023, March 27). How is India adapting to heatwaves an assessment of heat action plans with insights for transformative climate action. Centre for Policy Research.

Pörtner, H. O. (2004). Climate Variability and the Energetic Pathways of Evolution: The Origin of Endothermy in Mammals and Birds. https://Doi.Org/10.1086/423742, 77(6), 959–981. 10.1086/423742

Rekha, S., Nalini, S. J., Bhuvana, S., Kanmani, S., Hirst, J. E., & Venugopal, V. (2024). Heat stress and adverse pregnancy outcome: Prospective cohort study. BJOG : An International Journal of Obstetrics and Gynaecology, 131(5). 10.1111/1471-0528.17680

Riley, K., Wilhalme, H., Delp, L., & Eisenman, D. P. (2018). Mortality and Morbidity during Extreme Heat Events and Prevalence of Outdoor Work: An Analysis of Community-Level Data from Los Angeles County, California. International Journal of Environmental Research and Public Health, 15(4). 10.3390/IJERPH15040580

Roos, N., Kovats, S., Hajat, S., Filippi, V., Chersich, M., Luchters, S., Scorgie, F., Nakstad, B., Stephansson, O., Hess, J., Kadio, K., Kouanda, S., Lusambili, A., Marsham, J., Ngugi, A., & Wright, C. Y. (2021). Maternal and newborn health risks of climate change: A call for awareness and global action. Acta Obstetricia et Gynecologica Scandinavica, 100(4), 566–570. 10.1111/AOGS.14124

Sixth Assessment Report — IPCC. (n.d.). Retrieved April 4, 2024, from https://www.ipcc.ch/assessment-report/ar6/

Skibiel, A. L., Peñagaricano, F., Amorín, R., Ahmed, B. M., Dahl, G. E., & Laporta, J. (2018). In Utero Heat Stress Alters the Offspring Epigenome. Scientific Reports 2018 8:1, 8(1), 1–15. 10.1038/s41598-018-32975-1

Solano, M. E., Jago, C., Pincus, M. K., & Arck, P. C. (2011). Highway to health; or How prenatal factors determine disease risks in the later life of the offspring. Journal of Reproductive Immunology, 90(1), 3–8. 10.1016/J.JRI.2011.01.023

Strand, L. B., Barnett, A. G., & Tong, S. (2011). The influence of season and ambient temperature on birth outcomes: a review of the epidemiological literature. Environmental Research, 111(3), 451–462. 10.1016/J.ENVRES.2011.01.023

Sun, S., Weinberger, K. R., Spangler, K. R., Eliot, M. N., Braun, J. M., & Wellenius, G. A. (2019). Ambient temperature and preterm birth: A retrospective study of 32 million US singleton births. Environment International, 126, 7–13. 10.1016/J.ENVINT.2019.02.023

Takahashi, M. (2012). Heat stress on reproductive function and fertility in mammals. Reproductive Medicine and Biology, 11(1), 37. 10.1007/S12522-011-0105-6

Thornton, P., Nelson, G., Mayberry, D., & Herrero, M. (2022). Impacts of heat stress on global cattle production during the 21st century: a modelling study. The Lancet Planetary Health, 6(3), e192–e201. 10.1016/S2542-5196(22)00002-X

Walsh, K., McCormack, C. A., Webster, R., Pinto, A., Lee, S., Feng, T., Sloan Krakovsky, H., O’Grady, S. M., Tycko, B., Champagne, F. A., Werner, E. A., Liu, G., & Monk, C. (2019). Maternal prenatal stress phenotypes associate with fetal neurodevelopment and birth outcomes. Proceedings of the National Academy of Sciences of the United States of America, 116(48), 23996–24005. 10.1073/PNAS.1905890116/-/DCSUPPLEMENTAL

Wang, J., Williams, G., Guo, Y., Pan, X., & Tong, S. (2013). Maternal exposure to heatwave and preterm birth in Brisbane, Australia. BJOG : An International Journal of Obstetrics and Gynaecology, 120(13), 1631–1641. 10.1111/1471-0528.12397

Watkins, A. J., Ursell, E., Panton, R., Papenbrock, T., Hollis, L., Cunningham, C., Wilkins, A., Perry, V. H., Sheth, B., Wing, Y. K., Eckert, J. J., Wild, A. E., Hanson, M. A., Osmond, C., & Fleming, T. P. (2008). Adaptive Responses by Mouse Early Embryos to Maternal Diet Protect Fetal Growth but Predispose to Adult Onset Disease. Biology of Reproduction, 78(2), 299–306. 10.1095/BIOLREPROD.107.064220

Zhang, Y., Yu, C., & Wang, L. (2017). Temperature exposure during pregnancy and birth outcomes: An updated systematic review of epidemiological evidence. Environmental Pollution, 225, 700–712. 10.1016/j.envpol.2017.02.066

Zhao, W., Liu, F., Bell, A. W., Le, H. H., Cottrell, J. J., Leury, B. J., Green, M. P., & Dunshea, F. R. (2020). Controlled elevated temperatures during early-mid gestation cause placental insufficiency and implications for fetal growth in pregnant pigs. Scientific Reports, 10(1), 20677. 10.1038/s41598-020-77647-1

